# The delta-opioid receptor bidirectionally modulates itch

**DOI:** 10.1101/2022.08.09.503338

**Authors:** Kelly M. Smith, Eileen Nguyen, Sarah E Ross

**Affiliations:** University of Pittsburgh School of Medicine, Department of Neurobiology; University of Pittsburgh, Pittsburgh Center for Pain Research; University of Pittsburgh School of Medicine, Medical Scientist Training Program

**Keywords:** Itch, pruritus, DOR, delta opioid, spinal cord

## Abstract

Opioid signaling has been shown to be critically important in the neuromodulation of sensory circuits in the superficial spinal cord. Agonists of the mu-opioid receptor (MOR) elicit itch, whereas agonists of the kappa-opioid receptor (KOR) have been shown to inhibit itch. Despite the clear roles of MOR and KOR for the modulation itch, whether the delta-opioid receptor (DOR) is involved in the regulation of itch remained unknown. Here, we show that intrathecal administration of DOR agonists suppresses chemical itch and that intrathecal application of DOR antagonists is sufficient to evoke itch. We identify that spinal enkephalin neurons co-express neuropeptide Y (NPY), a peptide previously implicated in the inhibition of itch. In the spinal cord, DOR overlapped with both the NPY receptor (NPY1R) and KOR, suggesting that DOR neurons represent a site for convergent itch information in the dorsal horn. Lastly, we found that neurons co-expressing DOR and KOR showed significant Fos induction following pruritogen-evoked itch. These results uncover a role for DOR in the modulation of itch in the superficial dorsal horn.

**Perspective:** This article reveals the role of the delta opioid receptor in itch. Intrathecal administration of delta agonists suppresses itch whereas the administration of delta antagonists is sufficient to induce itch. These studies highlight the importance of delta-opioid signaling for the modulation of itch behaviors, which may represent new targets for the management of itch disorders.

## Introduction

Acute itch is an important sensation that alerts organisms to stimuli that need to be removed from the skin. Investigation into the cells and circuits involved in the transmission of itch has identified several targets both peripherally (Mrgpra3+, Mrgprd+ and Nppb/Sst+)^1-6^ and centrally (gastrin releasing peptide and its receptor and substance P and the neurokinin 1 receptor) that promote itch^7-13^.

Within the spinal cord, opioid peptides and their receptors have been found to be important for both pain and itch. In particular, the role of the mu- and kappa-opioid receptors have been well described. Agonism of the mu opioid receptor (MOR) induces itch^14,15^, whereas antagonism of MOR suppresses itch^16,17^. Moreover, the effects of mu agonists on itch have been shown to specifically act through MOR, as deletion of this receptor attenuates itch responses to mu agonists^15,18^. In contrast, agonism of the kappa opioid receptor (KOR) inhibits itch^19-24^. Both partial and selective agonists of KOR are presently used in clinical settings for the management of various forms of pruritus^25-27^.

Despite the known role of MOR and KOR in spinal modulation of itch, the delta opioid receptor (DOR) has not been studied in this context. The role for DOR agonism in the suppression of nociception has previously been identified^28-33^. Intrathecal administration of DOR agonists inhibits allodynia associated with neuropathic injury^34-36^. The underlying mechanism for this observation is thought to occur through somatostatin spinal neurons containing DOR, as agonism of DOR inhibits SOM+ neurons, a cell type that has previously been shown to decrease mechanical allodynia^37^. Interestingly in the context of itch, somatostatin neurons have been implicated in spinal itch pathways. In particular, ablation of SOM+ neurons in the spinal cord decreased scratching induced by common itch causing agents such as compound 48/80 and chloroquine^38^. Combined, these data suggest that the DOR may be involved in the modulation of spinal itch circuits.

Other neuropeptides and their receptors have also been implicated in the spinal modulation of itch. Several labs have shown that neuropeptide Y (NPY) and its receptor, NPY1R, are critical for mechanical itch. Spinal administration of a NPY1R agonist decreases mechanically evoked scratching^39^, while ablation of NPY1R expressing spinal neurons prevents mechanical itch^10^. Neurons that express dynorphin (Dyn), the endogenous ligand for KOR has been shown to inhibit chemical itch. In particular, mice that lack Dyn expressing neurons due to the loss of Bhlhb5 during development develop pathalogical itch^19,40^, whereas as chemogenetic activation of Dyn neurons in the spinal cord decreases chloroquine induced scratching^6^. These studies highlight that multiple peptide signaling pathways can modulate itch. However, we do not yet understand the degree to which these distinct peptides and their respective receptors are expressed on common (vs. distinct) populations of neurons, and this gap of knowledge precludes a clear picture of the logic of neuropeptide signaling for itch.

Other neuropeptides and their receptors have also been implicated in the spinal modulation of itch. Several labs have shown that neuropeptide Y (NPY) and its receptor, NPY1R, are critical for mechanical itch. Spinal administration of a NPY1R agonist decreases mechanically evoked scratching^39^, while ablation of NPY1R expressing spinal neurons prevents mechanical itch^10^. Neurons that express dynorphin (Dyn), the endogenous ligand for KOR has also been shown to inhibit chemical itch as chemogenetic activation of Dyn neurons in the spinal cord decreases chloroquine induced scratching^6^. These studies show that itch, at least in the spinal cord is modulated by both neuropeptides and opioids.

Here we show that agonism of DOR decreased scratching behavior, whereas antagonism of DOR generates spontaneous scratching. Further we show that enkephalin (Enk, the endogenous ligand for DOR) is co-expressed on NPY neurons and that KOR and DOR are often co-expressed on the same population. Taken together these data raise the possibility that NPY and Dyn neurons which release Enk and Dyn respectively, share a common neural target—one that expresses both KOR and DOR— to mediate the inhibition of itch.

## Materials and Methods

### Mice

Male and female C57Bl/6 mice aged 6-10 weeks (17-25 g) were used in these studies (Charles River Laboratories). Even numbers of male and female mice were used for all experiments and no clear sex differences were observed so data were pooled. Mice were housed under standard laboratory conditions with temperature and humidity-controlled environment and a 12 h light/dark cycle. Standard lab chow and water were available *ad libitum*. The use of animals was approved by the University of Pittsburgh Animal Care and Use Committee.

### Pharmacologic agents

Deltorphin II (Sigma), naltrindole (Sigma), and chloroquine diphosphate salt (Sigma) were dissolved in physiological saline. TIPP Ψ (Tocris) was dissolved in 10% DMSO. SNC80 (Tocris) was dissolved in 0.2% HCl saline. Deltorphin, naltrindole, TIPP, and SNC80 were administered intrathecally (doses provided in results and figure legends). Chloroquine was administered intradermally (described further below).

### Intradermal injection of chloroquine

For intradermal injection of chloroquine, hair was removed at least 24 h before the experiment. Chloroquine (100 μg in 10 μL) was administered intradermally into the nape of the neck, which could be visualized by the formation of a small bubble under the skin. To assess the inhibition of chloroquine-induced itch by delta opioid receptor agonists, chloroquine was injected 30 min following IT deltorphin II or SNC80.

### Intrathecal injections

For intrathecal injections, hair was clipped from the back of each mouse at least 24 h before the experiment. All intrathecal injections were delivered in a total volume of 10 μL using a 30-gauge needle attached to a luer-tip 25 μL Hamilton syringe. The needle was inserted into the tissue at a 45° angle and through the fifth intervertebral space (L5 – L6). Solution was injected at a rate of 1 μL/s. The needle was held in position for 10 s and removed slowly to avoid any outflow of the solution. Only mice that exhibited a reflexive flick of the tail following puncture of the dura were included in our behavioral analysis. These procedures were performed in awake mice.

### Behavior

All assays were performed in the Pittsburgh Phenotyping Core and scored by an experimenter blind to treatment.

### Observation of scratching behavior

Scratching behavior was quantified as described previously^19^. In brief, on the testing day, the mice were individually placed in the observation cage (12×9×14 cm) to permit acclimation for 30 min. Scratching behavior was videotaped for 30 min after administration of chloroquine or for 60 minutes after the administration of delta antagonists. The total numbers of scratch bouts by the hind paws at various body sites during this period were counted by a person who was blinded to treatment.

### RNAscope in-situ hybridization

Multiplex fluorescent in-situ hybridization (FISH) was performed according to the manufacturer’s instructions (Advanced Cell Diagnostics #320850). Briefly, 16 μm-thick fresh-frozen sections containing the lumbar enlargement of WT mouse spinal cord were fixed in 4% paraformaldehyde, dehydrated, treated with protease for 15 minutes, and hybridized with gene- and species-specific probes. Probes use in these experiments include; Mm-Penk-C1 (#318761), Mm-Oprd1-C2 (#427371-C2), Mm-Oprk1-C1 (#316111), Mm-Npy-C2 (#313321-C2), Mm-Fos-C3 (#498401-C3), Mm-Slc32a1-C3 (#319191), Mm-Slc17a6-C3 (#319171), and Mm-Mpy1r-C3 (#427021). DAPI (#320858) was used to visualize nuclei. 3-plex positive (#320881) and negative (#320871) control probes were tested.

### Image acquisition and quantification

Full-tissue thickness sections were imaged using either an Olympus BX53 fluorescent microscope with UPlanSApo 4x, 10x, or 20x objectives or a Nikon A1R confocal microscope with 20X or 60X objectives. All images were quantified and analyzed using ImageJ. To quantify images in RNAscope in-situ hybridization experiments, confocal images of tissue samples (3-4 dorsal horns per mouse, with n = 3-5 mice) were imaged and only cells whose nuclei were clearly visible by DAPI staining and exhibited punctate fluorescent signal were counted.

### Statistics

All statistical analyses were performed using GraphPad Prism 9. Values are presented as mean ± SEM. P values were determined by tests indicated in applicable figure legends. Sample sizes were based on pilot data and are similar to those typically used in the field.

## Results

### The delta-opioid receptor bidirectionally modulates itch

Previous work has shown that both MOR and KOR within the spinal cord dorsal horn are important for the transmission of itch information^15,19^. To address whether DOR also plays a role in the modulation of itch at the level of the spinal cord we used intrathecal administration of DOR agonists (Figs 1A and 1B). We found that the both agonists, deltorphin II (1 µg) and SNC80 (45 ng) significantly decreased the number of scratch bouts associated with intradermal chloroquine (153.4 ± 21.6 vs 47.3 ± 9.9, p=0.0004 and 110.6 ± 7.3 vs 45 ± 10.5, p=0.0002, n=9 or 8, Fig 1C). Further, intrathecal administration of the DOR antagonists TIPP Ψ (10 ng) and naltrindole (13 ng, Figs 1D and 1E) significantly increased the number of spontaneous scratch bouts (32.0 ± 3.3 vs 92.5 ± 13.3, p=0.0006, and 14.7 ± 3.4 vs 56.7 ± 13.2, p=0.009, n=8 or 7, Fig 1F). Together these results suggest that DOR in the spinal cord bidirectionally modulates itch.

**Figure 1.**
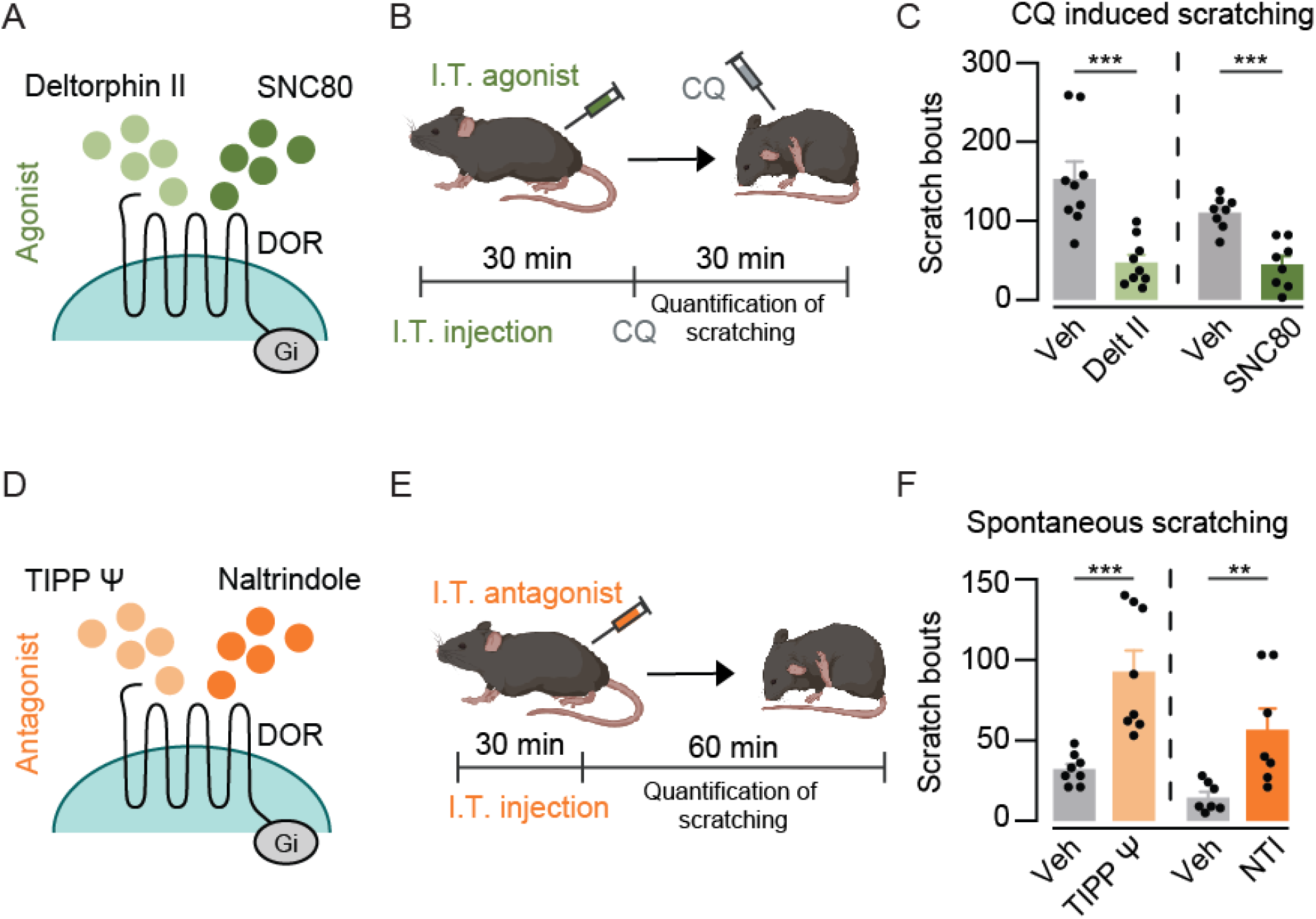
DOR bidirectionally modulates itch. **(A)** Schematic of DOR agonist action on neurons expressing the receptor. Both deltorphin II and SCN80 activate DOR, which is a Gi-coupled GPCR whose activation is predicted to decrease neuronal activity. **(B)** Schematic of experimental set up. WT mice were injected intrathecally (I.T.) with DOR agonists (deltorphin II (1 µg) or SNC80 (45 ng) or vehicle, left) and scratching behavior was quantified following intradermal chloroquine (CQ, 100 µg in 10 µL) injection to the nape of the neck (right). Timeline below shows when the compounds were administered. **(C)** Both deltorphin II (left, pale green) and SNC80 (right, dark green) significantly reduced the number of chloroquine induced scratch bouts compared to vehicle. Data are mean ± SEM with dots representing individual animals (n=8-9 mice), p=0.0004 (left), p=0.0002 (right), determined by students t-test. **(D)** Schematic of DOR antagonist action on neurons expressing DOR. Tipp Ψ and Naltrindole (NTI) inhibit DOR. **(E)** Schematic of experimental set up. WT mice were injected I.T. with DOR antagonists (TIPP Ψ (10 ng) or NTI (13 ng) or vehicle, left) and spontaneous scratching behavior was quantified (right). Timeline below shows when the compounds were administered. **(F)** The total number of spontaneous scratch bouts observed in 60 min was quantified. Both TIPP Ψ (left, light orange) and NTI (right, dark orange) significantly increased the number of spontaneous scratch bouts. Data are mean ± SEM with dots representing individual animals (n=7-9 mice), p=0.0006 (left), p=0.0093 (right), determined by students t-test.

### NPY+ neurons co-express enkephalin, the endogenous DOR agonist

Given that DOR bidirectionally modulates itch we next investigated neurons that may be upstream. In particular, we focused our efforts on neurons that produce Enk, the endogenous ligand for DOR. Single sequencing data suggest that enkephalin-expressing dorsal horn neurons also express NPY^41^. Importantly, NPY+ neurons have been shown to be important for the spinal inhibition of itch^10,39^. We therefore performed fluorescent in situ hybridization (FISH) for *Npy*, pro-enkephalin (*Penk*) and the inhibitory marker *Slc32a1* (Vgat), which labels GABAergic neurons (Fig 2A). Consistent with previous reports^42^, we found that 20.9 ± 2% of *Penk* neurons were inhibitory (based on *Slc32a1* expression, Fig 2B, n=5). Moreover, of inhibitory *Penk* neurons, the vast majority also expressed *Npy* (76.8 ± 2.3%, n=5) (Figs 2C and 2D). To follow up on this observation, we next examined the overlap between DOR and NPY1R, the endogenous receptors for Enk and NPY, respectively. We found that 39.6 ± 4.1% of *Oprd1* neurons also expressed *Npy1r* (Figs 2E and 2F, n=4). Together, our FISH analyses raise the possibility that inhibitory NPY+/Penk+ neurons are upstream of DOR+/NPY1R+ neurons and that their release of Enk, NPY, or a combination of both inhibits itch signaling (Fig 2G).

**Figure 2.**
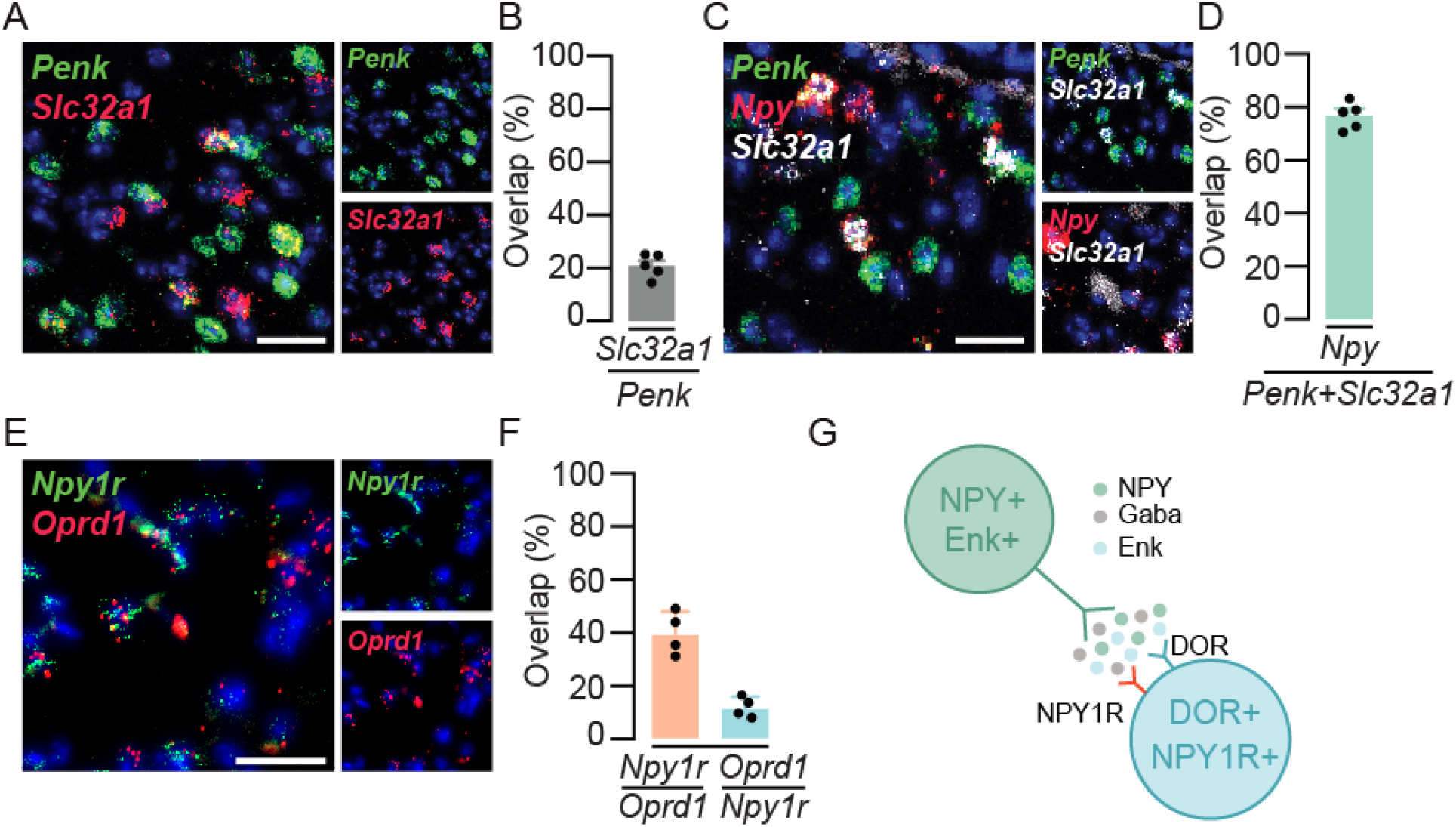
Inhibitory *Penk* neurons express *Npy*. **(A)** Representative image of FISH experiment for *Penk* (green), and *Slc32a1* (red). Scale bar = 25 µm. **(B)** Quantification of FISH labeled neurons in the SDH that express both *Slc32a1* and *Penk*. n = 3-4 sections from 5 animals, data are mean ± SEM with dots representing averaged data from individual animals. **(C)** Representative image of FISH experiment for *Penk* (green), *Npy* (red) and *Slc32a1* (white). Scale bar = 25 µm. **(D)** Quantification of FISH labeled *Slc32a1*+ neurons in the SDH that also express *Npy* and *Penk*. n = 3-4 sections from 5 animals. Data are mean ± SEM with dots representing averaged data from individual animals. **(E)** Representative image of FISH for *Npy1r* (green) and *Oprd1* (red). Scale bar = 25 µM. **(F)** Quantification of FISH labeled neurons in the SDH that express both *Npy1r* and *Oprd1*. n = 3-4 sections from 4 animals, data are mean ± SEM with dots representing averaged data from individual animals. **(G)** Predicted model for signaling between NPY+ and DOR+ neurons. NPY neurons are proposed to release both enkephalin and NPY onto neurons that are both DOR+ and NPY1R+.

### DOR and KOR are co-expressed in a population of dorsal horn neurons

KOR has a well-established role in the spinal inhibition of itch^15,19,25-27,43-46^. Therefore, we performed FISH to examine the co-expression of *Oprd1* and *Oprk1* on spinal neurons (Fig 3A). Quantification of FISH labelling showed ∼40% overlap between *Oprd1* and *Oprk1* in excitatory neurons marked by the expression of *Slc17a6*. Specifically, 38.7 ± 2.9% of *Slc17a6*+ and *Oprk1*+ neurons co-expressed *Oprd1* and 38 ± 4.1% of *Slc17a6*+ and *Oprd1+* neurons co-expressed *Oprk1* (Figs 3B and 3C, n=3). This overlap in expression of KOR and DOR, together with our previous finding that Penk is expressed in NPY neurons, suggested two possible models for DOR involvement in the inhibition of itch. In the first model, ‘parallel pathways’, NPY-expressing neurons are upstream of DOR neurons and Dyn neurons are upstream of KOR neurons (Fig 3D). This parallel model could explain the proposed divergent roles of NPY and Dyn in mechanical and chemical itch, respectively^10,19,39^. However, given the observed overlap of nearly half of KOR and DOR neurons, an alternative model, ‘convergent pathways’ was also a possibility. In this model, both NPY- and Dyn-expressing cells are upstream of a common neuron that co-expresses DOR and KOR (Fig 3E), indeed, there have been reports that NPY can also play a role in chemical itch^47,48^. If so, these DOR+/KOR+ neurons could represent a point of convergence for mechanical and chemical itch signaling.

**Figure 3.**
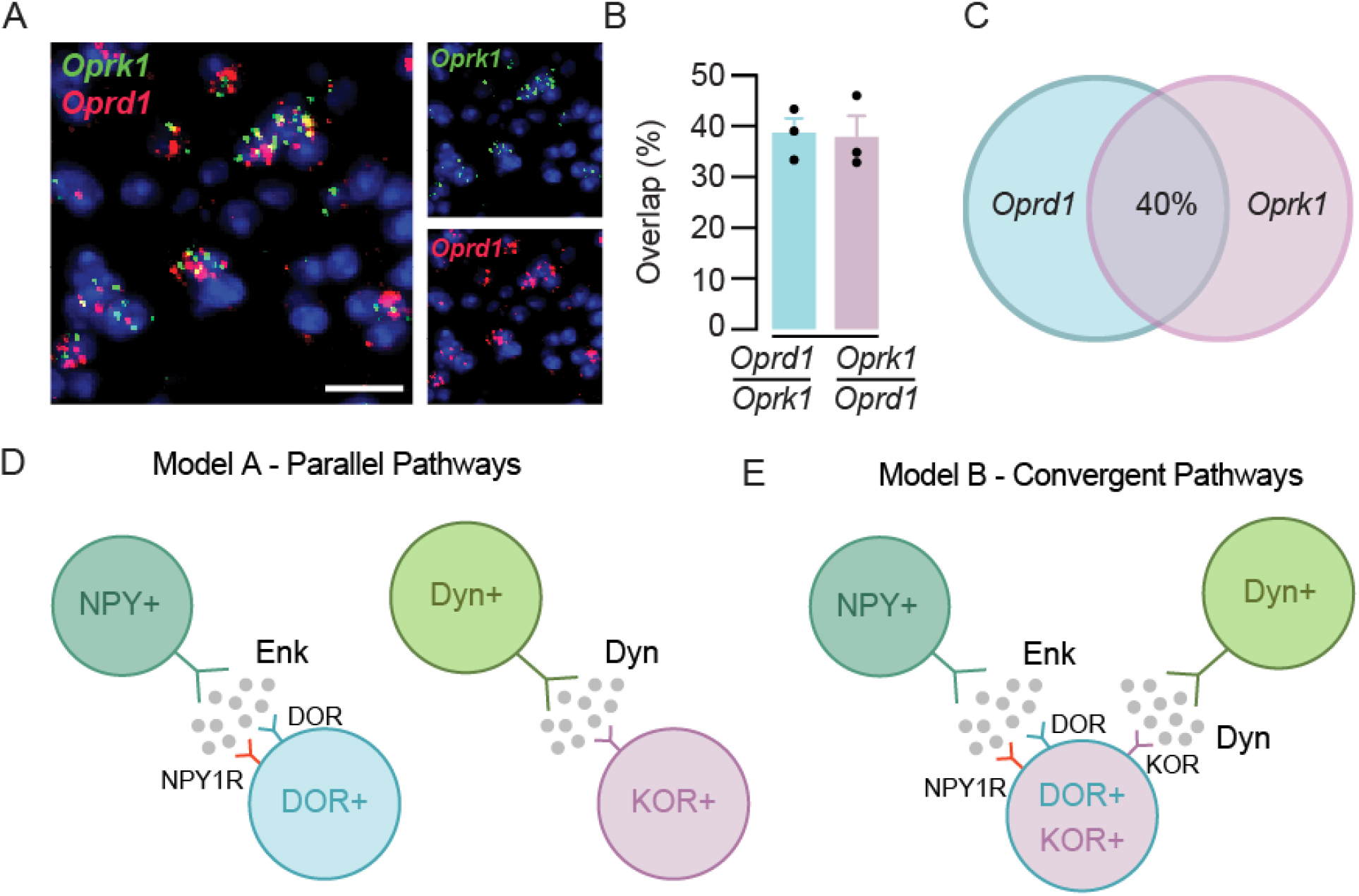
*Oprd1* and *Oprk1* are co-expressed in a population of excitatory neurons. **(A)** Representative image of FISH for *Oprd1* (red) and *Oprk1* (green). Scale bar = 10 µm. **(B and C)** Quantification of FISH labeled neurons in the SDH that express *Oprd1* and *Oprk1* in *Vglut2*+ cells, n = 3-4 sections from 3 animals. Data are mean ± SEM with dots representing averaged data from individual animals. **(D)** Model A schematic, Parallel Pathways. NPY+ neurons release enkaphalin (Enk) onto DOR+ neurons while Dyn neurons release dynorphin (Dyn) onto KOR+ neurons in separate circuits. **(E)** Model B schematic, Convergent Pathways. NPY+ neurons and Dyn neurons release Enk and Dyn, respectively, onto a population that co-expresses DOR and KOR.

### DOR and KOR neurons are activated following an itch stimulus

To investigate these potential models in more detail, we analyzed which neurons are activated in the context itch using Fos as a marker of neural activity. For these experiments, we used intradermal chloroquine to induce itch along with FISH (Figs 4A and 4B), using probes against *Oprk1, Oprd1* and *Fos*, to examine which population(s) were active. Neurons were classified based on their expression of opioid receptors as follows: *Oprk1* but not *Oprd1* (*Oprk1* alone), *Oprd1* but not *Oprk1* (*Oprd1* alone) or *Oprd1* and *Oprk1* co-expressing neurons (*Oprd1 + Oprk1*, Fig 4C). Quantification of FISH revealed that *Fos* was significantly induced in the *Oprk1 alone* population and the *Oprd1+Oprk1* populations. In particular, 18.7 ± 2.1% of *Oprk1 alone* neurons co-expressed *Fos* after chloroquine injection compared to 6.5 ± 2.3% following saline injection (p=0.047, n=3, Fig 4D). Similarly, a larger percentage of neurons that expressed both *Oprk1* and *Oprd1* co-expressed *Fos* following CQ administration (1 ± 2.4% vs 27.8 ± 3.7%, p=0.047, n=3, Fig 4D). No significant difference in co-expression with Fos was observed in *Oprd1* alone neurons (5.8 ± 2.9% vs 14.2 ± 0.8%, p=0.0502, n=3, Fig 4D), although the effect trended toward significance. Together, these data raise the possibility that the behavioral effects of DOR agonism and antagonism may occur, at least in part, through the action of DOR on neurons that co-express DOR and KOR. Furthermore, the observation that DOR+/KOR+ neurons are significantly activated by a pruritogen suggest that itch signaling could converge on the neurons that co-express both receptors.

**Figure 4.**
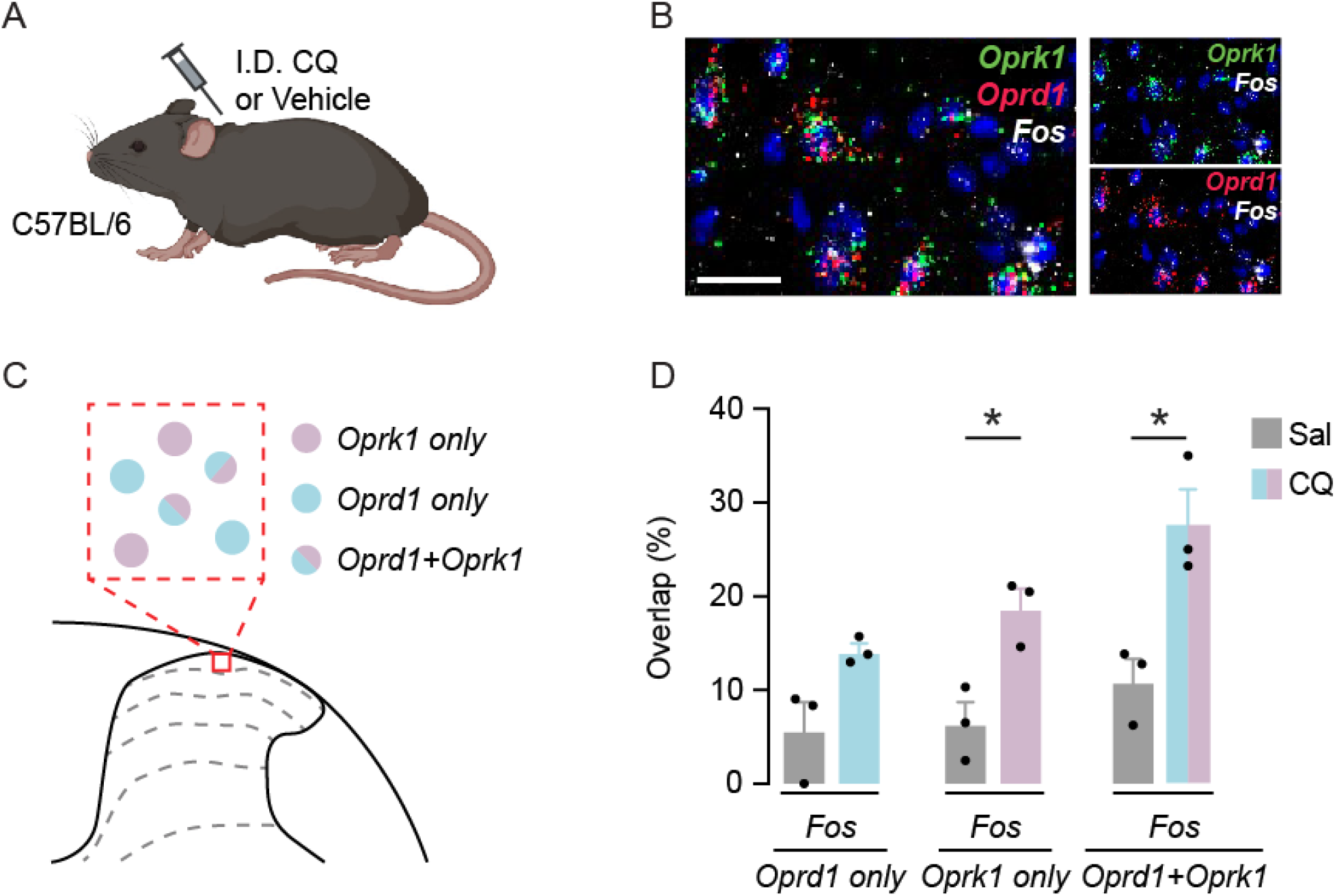
*Oprd1* and *Oprk1* co-expressing neurons are activated following an itch stimulus. **(A)** Schematic of experimental design. Wild type mice were injected intradermally (I.D.) with either chloroquine (100 μg in 10 μL) or vehicle in the nape of the neck. Spinal cords were subsequently processed for FISH. **(B)** Representative image of FISH for *Oprk1* (green), *Oprd1* (red) and *Fos* (white). Scale bar = 25 µm. **(C)** Schematic showing types of cells identified by FISH experiment, *Oprk1*+ only (mauve), *Oprd1*+ only (pale blue) and both *Oprk1*+ and *Oprd1*+ (split mauve and pale blue). **(D)** Quantification of FISH labelled neurons in the SDH that express *Fos* as well as *Oprd1* and/or *Oprk1*. n = 3-4 sections from 3 animals, data are mean ± SEM with dots representing averaged data from individual animals. P-values were determined by unpaired t-tests corrected for multiple comparisons using the Holm-Šídák method. P = 0.0471 (center), 0.0471 (right).

## Discussion

In this study, we expose the role of DOR in the bidirectional modulation of itch. We found that agonism of DOR suppresses chloroquine-induced scratching whereas antagonism of DOR enables spontaneous scratching. In addition, our studies revealed that inhibitory enkephalin neurons are enriched in their expression of NPY, a peptide that has previously been shown to be important for inhibition of mechanical itch^10,39^. We also found that DOR overlaps extensively with KOR, another opioid receptor that has been targeted successfully for the management of itch^25-27,43-45^. Finally, neurons containing both DOR and KOR showed significant Fos expression following chloroquine-induced itch. These DOR and KOR co-expressing neurons could represent the locus at which mechanical and chemical itch converge.

Several excitatory and inhibitory dorsal horn populations have previously been implicated in the spinal transmission of itch. Some of these populations have been reported to be specific for itch (GRP/GRPR,^11-13^), while others are likely to be involved in both nociception and itch (substance P/NK1R^7,49,50^). Spinal networks that act to inhibit itch have also been investigated, and these studies have implicated NPY/NPY1R pathways and kappa-opioid signaling pathways. In particular, our lab has previously shown that peripherally restricted kappa agonists reduce pain and itch behaviors^51^ and that agonism of KOR in the spinal cord reduces morphine induced itch^15^. Results from the current study extend on this previous work by revealing that, just like MOR and KOR, DOR also plays a modulatory role in the integration of itch at the level of the spinal cord.

One limitation of this study is that we were not able to test the role of spinal neurons containing DOR directly. Instead, we were able to manipulate these neurons using two different DOR agonists and two different DOR antagonists and found consistent results across the agents used. The reproducibility of our findings using distinct agonists supports the specificity of our agents for DOR. Furthermore, we applied our agents neuraxially, rather than systemically, to restrict the action of these agents directly on the neuraxis. It is important to note, however, that DOR is expressed on a population of large diameter dorsal root ganglia neurons and activity of these afferents could have been affected by intrathecal delivery of DOR agonists/antagonists. However, because DOR+ primary afferents are predicted to be involved in pain, rather than itch^30,52,53^, we favor the possibility that the effects of DOR manipulation observed here were do the its modulatory effect on spinal neurons rather than primary afferents.

We also cannot exclude the possibility that our intrathecal injection could have spread to the brain. However, the time frame within which we performed our experiments, which were conducted within an hour of intrathecal administration to reduce spread, may help to mitigate this concern. We also examined the expression of *Oprd1* in the spinal cord following of intradermal chloroquine, an itch-specific stimulus, and found robust expression of *Fos* in spinal neurons. These data are thus consistent with the possibility that the itch behaviors that were modulated by DOR signaling were due to its effects on spinal neurons rather than those in the brain. The future use of DOR-Cre mice, for instance, could permit the direct manipulation, recording, and imaging of DOR+ neurons in the spinal cord in the setting of itch to test this inference more directly.

Opioid receptors have been extensively investigated as targets for pain, and more recently, itch. Agonism of MOR, KOR, and DOR have all shown promise for their anti-nociceptive effects. However, their roles in itch are much more complex. It is intriguing that MOR and KOR have previously been shown to exhibit dichotomous roles – agonism of MOR elicits itch, whereas agonist of KOR suppresses itch. In this regard, DOR appears to be similar to KOR. Given that KOR agonists are used clinically to relieve itch, it may be that DOR will likewise be a useful pharmacological target. Thus, our discovery of the role of DOR in the modulation of itch may expand avenues for the management of itch.

